# cellstruct: Metrics scores to quantify the biological preservation between two embeddings

**DOI:** 10.1101/2023.11.13.566337

**Authors:** Jui Wan Loh, John F. Ouyang

**Author notes:** **Address for correspondence and reprint requests:** Dr. John F. Ouyang, PhD, Centre for Computational Biology; Programme in Cardiovascular and Metabolic Disorders, Duke-NUS Medical School, 8 College Rd, Singapore 169857.

## Abstract

Single-cell transcriptomics (scRNA-seq) is extensively applied in uncovering biological heterogeneity. There are different dimensionality reduction techniques, but it is unclear which method works best in preserving biological information when creating a two-dimensional embedding. Therefore, we implemented cellstruct, which calculates three metrics scores to quantify the global or local biological similarity between a two-dimensional and its corresponding higher-dimensional PCA embeddings at either single-cell or cluster level. These scores pinpoint cell populations with low biological information preservation, in addition to visualizing the cell-cell or cluster-cluster relationships in the PCA embedding. Two study cases illustrate the usefulness of cellstruct in exploratory data analysis.

## Background

Single-cell transcriptomics (scRNA-seq) is increasingly used to interrogate the biological heterogeneity and disease progression in different complicated biological systems. Various dimensionality reduction methods [1–3] are employed to reduce the high-dimensional features i.e. highly variable genes to two dimensions. t-SNE [4, 5] and UMAP [6, 7] are commonly used in single-cell analysis packages (e.g. Seurat [8] and Scanpy [9]) for removing technical noise while maintaining the biological signals both globally and locally. These two-dimensional embeddings evaluate the performance of multi-modal/batch integration, define cluster relationship including cell type annotation, present gene expression changes between different conditions, and infer trajectories driving developmental or disease progression [10–15], inherently assuming the preservation of biological information from the underlying gene expression space. However, dimension reduction often results in information loss because it is difficult to represent all the complex biological variation in 2D. Moreover, dimension reduction involves highly non-linear mathematical operations, introducing different degree of transformation to different parts of data. Therefore, the assessment of local and global structures preservation in the reduced embeddings from the untransformed data is critical for the selection of most accurate representation of the underlying variance for making rigorous biological inferences [1, 16–19], minimizing the chances of data misinterpretations caused by distortions introduced in dimension reduction [18, 20–22].

We present cellstruct to evaluate a reduced embedding’s ability to retain global or local relationships as compared to the reference embedding, at both single-cell and cluster level. Cell populations with low metric scores indicate poor biological information retention, guiding users to subset certain cell populations for closer inspection, or to tune the dimension reduction hyperparameters for the generation of new embeddings with better structure preservation. Thus, cellstruct is indispensable in scRNA-seq exploratory analysis, by assessing the fidelity of different two-dimensional embeddings and by revealing the underlying biological distances/relationships between cells or clusters.

## Results and Discussion

Cellstruct implements three metrics for assessing the preservation of global or local relationships between a reduced (e.g. UMAP) and a reference (e.g. PCA) embeddings at both cluster and single-cell level. It also provides heatmaps and dimension reductions for visualizing the pairwise cell/cluster distances in different embeddings (Figure 1A). We discovered that the metric assessing local relationships in the preservation of the k-nearest neighbors does not contribute to better interpretation of datasets, particularly in refining ambiguous/mixed cell types (detailed analysis in supplementary text; Figure S1). Thus, we focus on the preservation of global relationships at both single-cell and cluster level. To achieve this, we devised a global single-cell (GS) score quantifying the correlation between the global position of a single cell in the reduced and reference embeddings. Here, the global position of a single cell is given by its distance against a fixed number of randomly selected waypoint cells. Similarly, we devised a global cluster (GC) score by correlating the cluster-cluster distances.

**Figure 1.**
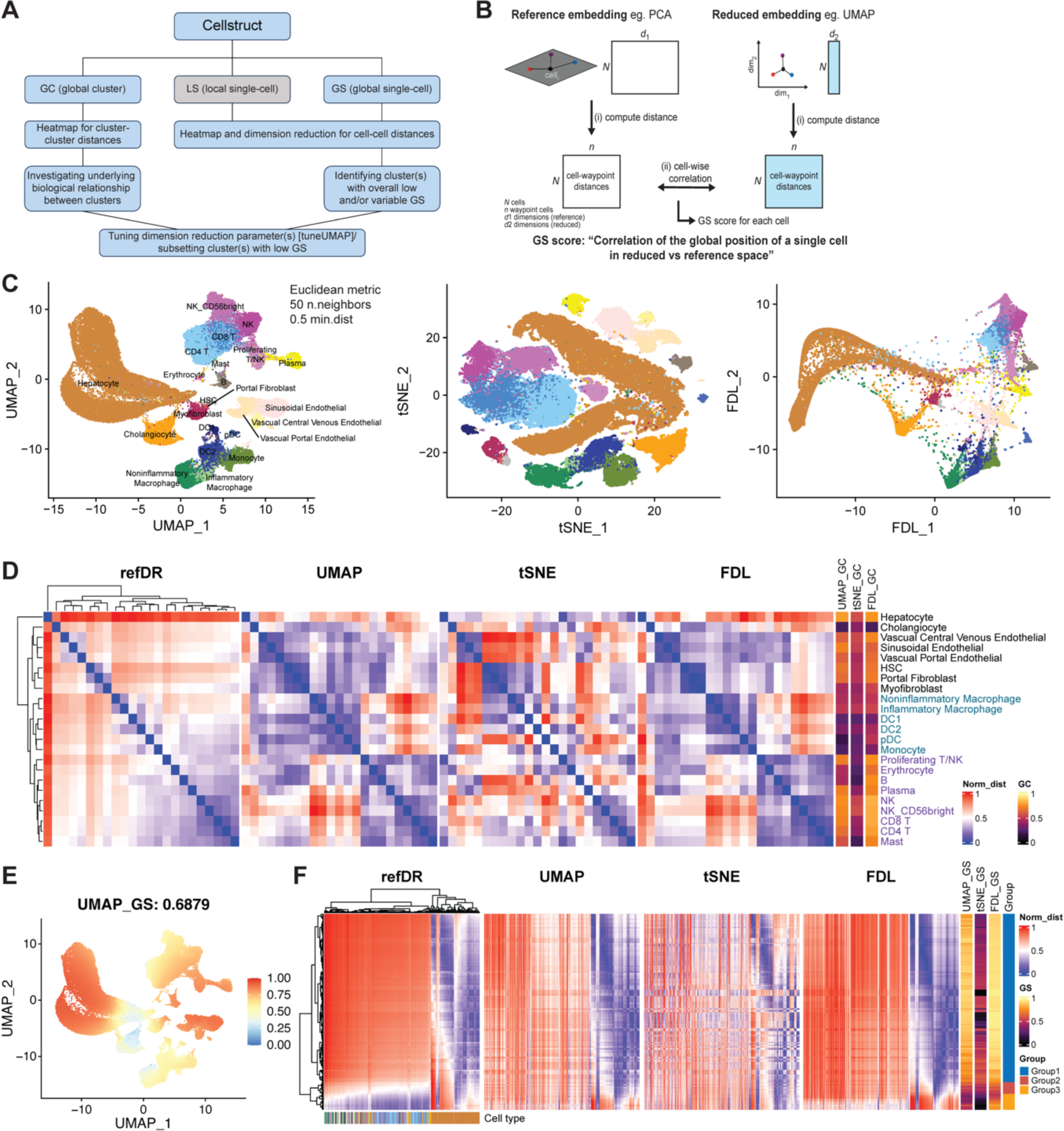
The cellstruct package and application on human liver cell atlas. (A) Schematic of cellstruct package, which includes three metrics scores and various visualizations to interrogate cell-cell or cluster-cluster relationships in single-cell data. (B) Schematic showing the calculation of global single-cell (GS) metric. (C) Cell type annotation of the liver cell atlas in UMAP, t-SNE, and FDL embeddings. (D) Heatmaps illustrating the normalized cluster-cluster distances in reference (i.e. refDR) and reduced (UMAP, t-SNE, and FDL) embeddings. GC scores of each cluster were depicted as single-column heatmaps for each reduced embedding, and cluster labels were colored by major grouping (blue: myeloid and purple: lymphoid and mast cell). (E) The distribution of GS scores on UMAP projection, with the mean score indicated in the title. (F) Heatmaps showing the cell-cell distances between 284 randomly sampled hepatocytes and 1,000 waypoint cells, in different embeddings. Again, GS scores were colored in the single-column heatmaps for each embedding. Three groups of hepatocytes were identified and visualized in each reduced embedding in Figure S5.

To illustrate these metrics, we applied cellstruct to a human liver cell atlas [23–27] (Figures 1A-C) where we calculated the GS and GC scores for different embeddings i.e. UMAP, t-SNE and Force-Directed Layout (FDL) (Figure S2). Note that we varied the hyperparameters to generate a tuned UMAP embedding, which will be used for the remaining analysis (further details in supplementary text; Figure S3). The cluster-cluster distance heatmap, which is used to calculate the GC score, revealed that hepatocytes are highly dissimilar from other cell types in the reference embedding (refDR), but this is not accurately reflected in reduced embeddings, particularly in t-SNE (Figure 1D). The lymphoid cells (B/T/NK cells, purple text in Figure 1D) were clustered closely with mast cells in all embeddings, resulting in high GC scores in general. Intuitively, one would expect the myeloid cells (macrophages/DCs, blue text) to be transcriptomically similar to the lymphoid cells, as correctly reflected in the refDR. However, the UMAP and FDL embeddings placed the myeloid closer to fibroblast/endothelial cells instead, indicated by the shorter distances in the cluster-cluster distance heatmap. Overall, all reduced embeddings recapitulated the relationships within the “islands” of endothelial and fibroblasts respectively, but t-SNE failed to reproduce it in the myeloid and lymphoid groups. The distance heatmap for t-SNE also less resembled the refDR counterpart, resulting in the lowest mean GC score amongst the different reduced embeddings (Figures 1D, S2).

At the single-cell level, we observed that the endothelial, lymphoid, and mast cells showed high GS scores, preserving the reference DR’s structure in the UMAP embedding (Figure 1E), and hepatocytes have the highest variance in GS score among all cell types (Figure S4), suggesting substantial heterogeneity in the refDR and/or UMAP embeddings. To investigate this heterogeneity, we sampled 284 hepatocytes (1%) to visualize the normalized pairwise distances between these cells and 1,000 randomly selected waypoint cells. Three distinct groups were observed to have very varied GS scores for all reduced embeddings. This variability in the GS score is alleviated using the FDL embedding, supported by the higher GS scores, particularly in Group3, due to the elongated projection of hepatocytes in FDL, which provided a better separation between Group1 and Group3 cells (Figure S5). Surprisingly, based on the cell-waypoint distance heatmap, Group2 and Group3 cells are transcriptomically more similar to other cell types, than the remaining hepatocytes (Figures 1F, S5).

We also interrogated the cholangiocytes due to their low GS scores. We sampled 428 cholangiocytes (∼10%) and identified 81 “outliers” (Group1 and Group 2 cells) with distinct cell-waypoint distance patterns (Figure S6). Three groups of cells were identified: Group1 cells are very likely mislabeled as hepatocytes, since they clustered together with the hepatocytes, Group2 cells represent a rare subpopulation, while Group3 cells are the main cholangiocyte population. Also, we noticed that FDL provided the worst embedding for cholangiocytes, particularly for Group3 cells, suggesting that different embeddings might be needed when interpreting different cell types (Figure S2). Overall, the FDL embedding is best at preserving the underlying global cell-cell and cluster-cluster relationships (further analysis in supplementary text; Figure S3).

<u>We next applied cellstruct to peripheral blood single-cell data from COVID-19 patients and healthy controls</u> [28]. The T/NK cells showed relatively high GS scores, while the remaining cell types have relatively lower GS scores, with COVID-CD14 Monocytes showing the most variability in GS score (Figures 2A, S8A). Thus, we decided to investigate the COVID-CD14 Monocytes further, sampled 1,657 cells (20%) for which we plotted the cell-waypoint distance heatmaps (Figures 2B-C, S8B). A small population of CD14 Monocytes did not cluster with the remaining cells of its type, and this small population comprises of only COVID-CD14 Monocytes, suggesting that these cells (black boxed) are different from other COVID-CD14 Monocytes (Figure S8B), driving the differences within these 20% sampled cells. Four groups of cells were identified, and they were well delineated in the tuned UMAP embedding of COVID-CD14 Monocytes only (Figures 2C-D). We revealed that these groups are significantly associated with the patient severity (Floor.NonVent, ICU.NonVent, and ICU.Vent) (*p*<2.2×10^−16^, Table S1), and Groups A-C shared similar gene set enrichment with more severe patients (GroupA-ICU.Vent: neutrophil degranulation, GroupC-ICU.Nonvent: interferon [IFN] signaling, GroupB: both biological processes) (Figures 2D-G). In addition, the monocytes analysis from Wilk et al. corroborated with our findings that very few cells within patients C2(1), C3(1), and C7(0) are found in Groups B and C (Table S2), as these patients have an absence of predicted IFN and IFN regulatory factor activities (Figure S9).

**Figure 2.**
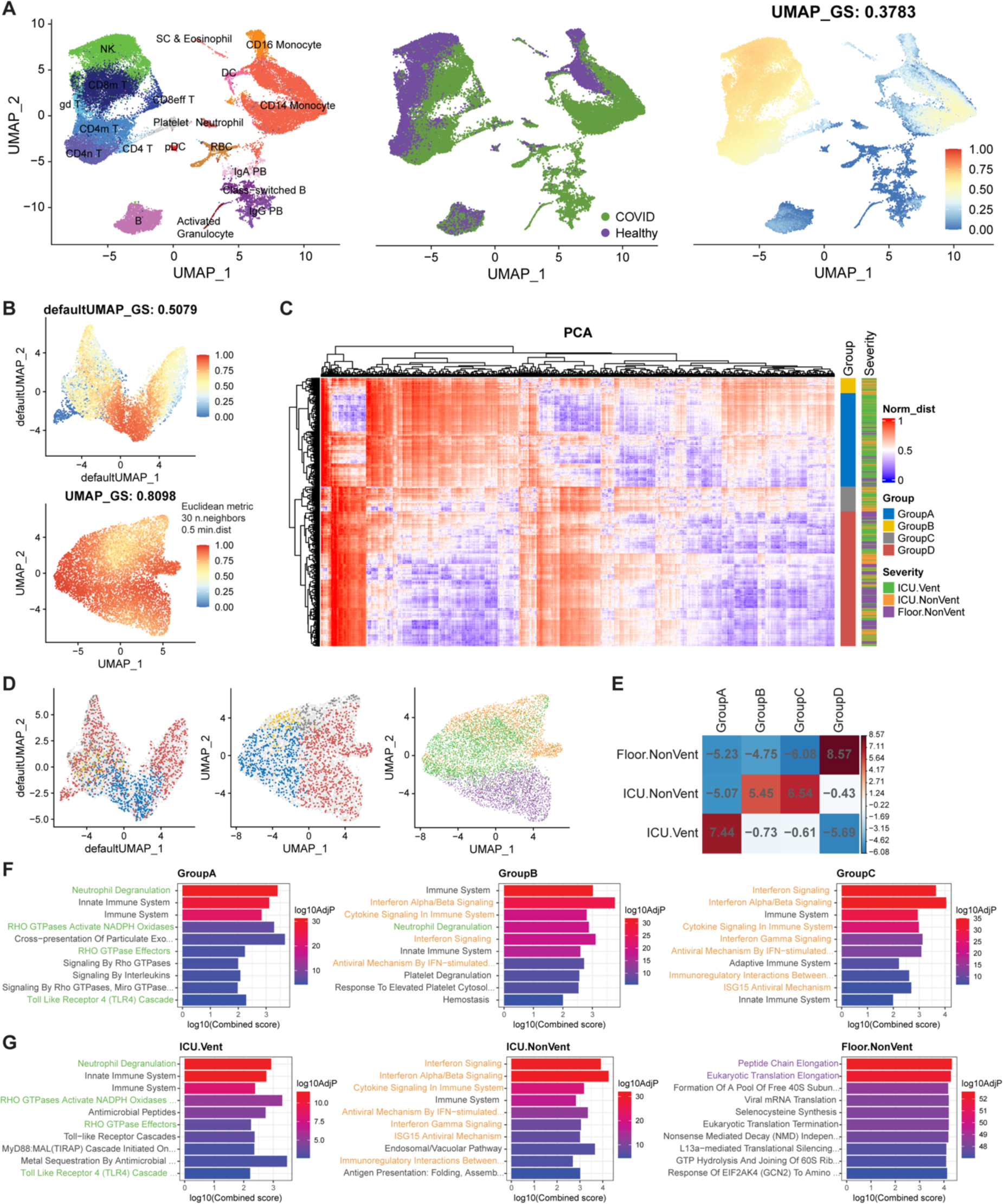
Application of cellstruct to the peripheral immune atlas of healthy and COVID patients. (A) UMAP embeddings showing the cell type annotation, COVID/healthy samples, and GS scores (mean score in the title) of the immune atlas. (B-G) The downstream analysis was focused on the 8,285 COVID-CD14 Monocytes subset, due to their high variability in GS score. (B) GS scores of COVID-CD14 Monocytes shown on default and tuned UMAP embeddings. (C) Heatmap showing cell-cell distances between 1,657 randomly sampled COVID-CD14 Monocytes (same cells as Figure S8B) and 1,000 waypoint cells in reference PCA embedding, divided into four groups of cells and annotated patient severity. (D) These four groups were delineated on default and tuned UMAP embeddings, and patient severity was only shown on the tuned UMAP, which does not show a separation of GroupD cells. (E) A contingency table of the four monocyte groups and patient severity, colored by the residuals of Chi-squared test. (F-G) Enriched pathways in different monocyte groups (F) and patient severity (G), taken from up-regulated genes between the group/severity of interest against remaining cells. Here, the randomly sampled COVID-CD14 Monocytes from Figure 2C were used, and GroupD monocytes were omitted due to small number of up-regulated genes. Groups A-C are mainly enriched for neutrophil degranulation and interferon signaling processes, which are respectively detected in ICU.Vent and ICU.NonVent cells. Floor.Nonvent cells are enriched for translation process. Pathways specific to patient severity were colored green, orange and purple.

Finally, we compared cellstruct with scDEED using their 20 simulated datasets (detailed comparison of simulated dataset 1 in supplementary text; Figure S10). scDEED assesses the reliability of reduced embeddings and classifies the cells into trustworthy and dubious [29]. We employed the same statistical approach to classify trustworthy cells using our GS score. With the respective set of trustworthy cells separately determined by cellstruct and scDEED, we measured the preservation of neighboring information using K-nearest neighbors (KNN) and K-nearest clusters (KNC) metrics. Cellstruct showed significantly higher KNN values in both t-SNE and UMAP embeddings (mean difference ≥0.15, *p*<1×10^−3^), while scDEED exhibited a higher KNC value in t-SNE (mean difference: 0.17, *p*=0.01) (Figure S11), suggesting that the trustworthy cells identified by cellstruct are more robustly preserved in the cell-cell relationships than the scDEED counterpart. Moreover, the computational time for cellstruct is one-third shorter than scDEED (cellstruct: ≤20 seconds, scDEED: ≤60 seconds) for both t-SNE and UMAP embeddings. Similar to scDEED, our tool is applicable to different embedding methods, and only the embeddings (and cell annotation for GC score) are required to run cellstruct.

## Conclusions

In this work, we demonstrated the utility of cellstruct for exploratory data analysis given different reduced embeddings. We provided several metrics and visualizations, which quantify the biological information preservation in a reduced embedding, facilitating the process in making observation-based biological interpretation. Users can now evaluate cell-cell or cluster-cluster similarity in the underlying high-dimensional space, and they are cautioned with cell populations comprising low/variable metric scores, avoiding potential misinterpretations from the highly non-linear dimension reduction procedure. More importantly, we hope cellstruct can bring awareness to the single-cell analysis community that while 2D embeddings are useful for visualization and interpretation, such embeddings often transform the underlying data to different extent for different cell populations.

## Methods

### Cellstruct

Cellstruct implements three metrics scores to quantify the preservation of global or local relationships between a reduced and a reference embedding at both global and local level. Each cell is assigned with GS (global single-cell) and optionally LS (local single-cell) scores that measure the correlation of its global position between both embeddings (Figure 1B) and the distances of its nearest neighbors within each embedding respectively (Figure S1A). Similarly, each cluster (i.e. cell type) is assigned with a GC (global cluster) score to describe the preservation of cluster-cluster relationship.

#### Global single-cell (GS) metric score

GS score is the Pearson (default) cell-wise correlation of the cell-waypoint distances calculated in the reference and reduced embeddings, given by this formula for cell *i*:

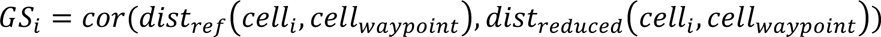

where 10,000 cells are randomly selected as “waypoint” cells that serve to describe the global position of each single cell, and pairwise distances between a single cell and these waypoints are computed in both reference and reduced embeddings to give the cell-waypoint distances. Note that the same set of waypoints is used across all single cells. Thus, the GS score indicates how well the global location of each cell is preserved from the reference to reduced embeddings.

#### Local single-cell (LS) metric score

LS score is the ratio of the mean reciprocal-squared-distance of the 30 nearest neighbors (NN) in the reduced embedding to the 30NN in the reference embedding, given by this formula for cell *i*:

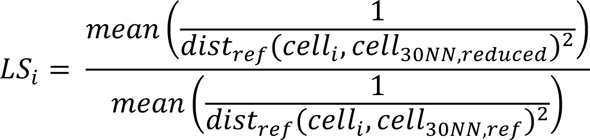

where the two sets of 30NN of the single cell (i.e. target) are determined using RANN R package (v.2.6.1), in the reduced and reference embeddings respectively. Thus, the LS score measures how far the reduced embedding NN are in the reference space, and the denominator serves as a normalization factor, so that LS varies from 0 to 1.

#### Global cluster (GC) metric score

GC score is the Pearson (default) correlation between the cluster-cluster distances calculated in the reference and reduced embeddings. Similar to the GS score, GC score evaluates how well the global location of a cluster is preserved between reference and reducing embeddings. Here, the distances are computed between the centroid of each cluster. The centroids are determined by averaging each dimension across all the cells in a cluster.

Overall, cellstruct takes in a Seurat object and requires users to specify the two input embeddings (i.e. the reduced and reference embeddings) and optionally cell annotations (e.g. cell type) for the GC score calculation. If cell annotations were not provided, Seurat clusterID (i.e. seurat_clusters) will be used instead, giving rise to less biological interpretable scores. For example, no biological inference could be made with the observation of short distance between the arbitrarily labeled clusters 1 and 2, as compared to the more meaningful labels e.g. CD4 and CD8 T cells. By default, cellstruct will calculate both the GC and GS scores and add these scores into the metadata of the returned Seurat object. In addition, dimension reduction plots illustrating the distribution of GC and GS scores as well as the cluster annotation will be generated for each reduced embedding (Figure S2).

Heatmaps and dimension reduction plots are provided as functions to illustrate the normalized pairwise distances in different embeddings, for better visualization of the metrics scores. For each cell (or each row in the cell-waypoint distance heatmap), the cell-cell distances are normalized by the 99 percentile of the distances between the single cell of interest and the remaining cells (or waypoint cells) in each embedding. For the purpose of illustration, 1,000 randomly selected cells are used in plotting heatmap. As for the GC metric visualization (i.e. the cluster-cluster distance heatmap), cluster-cluster distances are normalized by the maximum distance in each embedding.

### Datasets

Two single-cell datasets (human liver cell atlas from Azimuth [https://zenodo.org/record/7770308] and peripheral immune atlas from Fred Hutch [https://atlas.fredhutch.org/fredhutch/covid/dataset/wilk]) were used to demonstrate our tool. These datasets were preprocessed by the authors, and the Seurat objects were downloaded for our study.

Human liver cell atlas consists of 79,492 cells, which are composed of 23 cell types from the broader groups of hepatocytes, cholangiocytes, fibroblasts, immune cells (myeloid, T/NK, B, plasma cells, and erythrocytes), mast, endothelial, and stem cells (Figure 1C). This reference atlas was collated from several liver studies [23–27], involving 29 donors across a range of ages, for better liver cells annotation and understanding of the liver-related diseases. We generated UMAP, t-SNE, and FDL embeddings using the default hyperparameters, and as discussed in the supplementary text, we also selected another UMAP embedding with different hyperparameters (Euclidean metric, 50 n.neighbors, and 0.5 min.dist), which has the highest mean GS score for the exploratory analysis. For the investigation on cholangiocytes, we identified 78 “outliers” from all 3,634 cells via the clustering pattern detected in the cell-waypoint reference distance heatmap (data not shown), and these 78 “outliers”, along with the 350 randomly chosen non-repeating cholangiocytes, were used to plot the cell-waypoint distance heatmaps in Figure S6A.

This peripheral immune atlas was generated by Wilk et al. group to study the pathophysiology of COVID-19. It consists of 44,172 cells, comprising of T/NK, myeloid, B, red blood cells, plasmablasts, platelets, and granulocytes. The peripheral blood mononuclear cells (PBMCs) were collected from seven hospitalized COVID-19 patients (four of them developed acute respiratory distress syndrome) and six controls [28] (Figure 2A left, middle). For this study case, we focused on the COVID-19 CD14 Monocytes, as explained in the Results and Discussion section. 8,285 COVID-CD14 Monocytes were extracted, and UMAP embedding was tuned due to non-uniform distribution of GS scores across the same cell population and the detection of a group of partly separated cells with low GS scores on default UMAP (tuned UMAP parameters: Euclidean metric, 30 n.neighbors, and 0.5 min.dist) (Figure 2B). Pearson’s Chi-squared test was used to evaluate the association between the four detected groups of COVID-CD14 Monocytes and patient severity within 1,657 cells (Table S1, Figure 2E).

### Generation of UMAP, t-SNE, and FDL embeddings

Both UMAP and t-SNE embeddings were implemented with the functions RunTSNE() and RunUMAP() in the Seurat package. The hyperparameters that we tuned for UMAP embedding are metric, n.neighbors, and min.dist in the RunUMAP() function. The default hyperparameters of RunUMAP() and RunTSNE() are respectively cosine metric; 30 n.neighbors; 0.3 min.dist and 30 perplexity value. As for FDL, it was generated using scanpy.tl.draw_graph function in the Scanpy package. All the arguments were kept as default, unless indicated as otherwise, when we ran these functions.

### Stability analysis of cellstruct

We performed a stability analysis on the number of waypoint cells for GS score calculation, by varying the number of waypoints from 1K to 10K, with 1K increment on UMAP, t-SNE, and FDL embeddings of human liver cell atlas. The GS scores are independent of the number of waypoint cells from 1K to 10K cells, regardless of the dimension reduction methods (Figure S7A). Hence, we use 10K cells for GS score computation and 1K cells for heatmap illustration. In addition, we inspected the stability of tuneUMAP function, by downsampling the number of cells in human liver cell atlas to 5K, 8K, 10K, 20K, …, 60K, and 70K cells respectively, studying the performance of GS score across different dataset size. It was shown that GS scores are relatively consistent, with at least 20K cells being sampled (Figure S7B).

### Comparison with scDEED

scDEED (v0.1.0) is implemented as an R package. It provides a reliability score, which is the Pearson correlation of the target’s distances to its closest 50% neighboring cells in both reference and reduced embeddings, and a classification of dubious and trustworthy cells, by comparing to a null distribution of reliability score [29]. To compare cellstruct with scDEED, the same dubious/trustworthy cell classification, implemented by scDEED, was performed by comparing our GS score to a null distribution, which is the GS score assigned to a permuted object generated by scDEED. Using the default t-SNE and UMAP embeddings from the 20 simulated datasets generated by the scDEED authors, the preservation of neighboring information was evaluated using the same two metrics (K-nearest neighbors, KNN and K-nearest clusters, KNC) employed by them. Trustworthy cells, which were separately identified by cellstruct and scDEED in the t-SNE and UMAP, in the simulated datasets were retained (Table S3). Default t-SNE and UMAP embeddings were regenerated for these trustworthy cells, and dubious/trustworthy cells were re-classified for the evaluation of KNN and KNC metrics, assessing the biological preservation and also the robustness of each tool in identifying trustworthy cells (i.e. the first round of trustworthy cells). Paired t-test was used to statistically evaluate the differences in mean KNN and KNC values (Figure S11).

### Gene set enrichment analysis

Differential expression analysis was performed among the four monocyte groups, identified by cellstruct, and the three patient severity groups respectively from the 20% COVID-CD14 Monocytes subset using FindAllMarker function in Seurat (v4.3.0). The differential genes were tabulated in Tables S4 (monocyte groups) and S5 (patient severity) respectively. Since Group D has only seven up-regulated genes (adjusted *p*<0.05), gene set enrichment analysis was performed in Groups A-C and the three patient severity groups using enrichR (v3.2) [30–32] and Reactome 2022 database [33]. The top 10 significant biological processes were shown on the bar plots in Figures 2F-G.

## Availability of data and materials

The cellstruct R package can be installed from https://github.com/the-ouyang-lab/cellstruct. Data objects and codes for reproducing the figures and analysis can be found at https://github.com/the-ouyang-lab/cellstruct-reproducibility.

## Acknowledgements

Not applicable.

## Authors’ contributions

JWL and JFO wrote and edited the manuscript. JWL implemented the tool. JFO supervised the work.

## Ethics approval and consent to participate

Not applicable.

## Consent for publication

Not applicable.

## Supplementary Information

Additional file 1: Supplementary text (detailed discussions of LS metric and tuning UMAP hyperparameters, as well as comparative analysis of reduced embeddings and cellstruct vs scDEED)

Additional file 2: Supplementary figures (**Figure S1.** Evaluation of local cell-cell relationships via local single-cell (LS) metric in human liver cell atlas. **Figure S2.** An example of cellstruct output using the human liver cell atlas. **Figure S3.** Four single cells were selected as an illustration for the comparison between different reduced embeddings. **Figure S4**. Distribution of GS scores for UMAP embedding in human liver cell atlas. **Figure S5.** Dimension reductions showing three groups of hepatocytes. **Figure S6**. Investigating cholangiocytes in the human liver cell atlas using cellstruct. **Figure S7.** Stability analysis of GS metric using human liver cell atlas. **Figure S8.** Applying cellstruct to the COVID peripheral immune atlas. **Figure S9**. Corroboration of COVID-CD14 Monocyte groups, identified by cellstruct, with the author’s original analysis. **Figure S10.** Comparative analysis between scDEED and cellstruct using Simulated Dataset 1. **Figure S11.** Scatterplot of KNC and KNN values of t-SNE and UMAP embeddings in 20 simulated datasets.)

Additional file 3: Table S1. Number of cells for each group detected in different patient severity level.

Additional file 4: Table S2. Number of cells for each sample in each COVID-CD14 Monocyte group.

Additional file 5: Table S3. Number of trustworthy, dubious, and neither cells classified by scDEED and cellstruct respectively.

Additional file 6: Table S4. Differential genes expression analysis among four monocyte groups detected in 20% of COVID-CD14 Monocytes.

Additional file 7: Table S5. Differential genes expression analysis among three patient severity groups detected in 20% of COVID-CD14 Monocytes.

## References

1. Sun S, Zhu J, Ma Y, Zhou X: Accuracy, robustness and scalability of dimensionality reduction methods for single-cell RNA-seq analysis. Genome Biol 2019, 20:269.

2. Xiang R, Wang W, Yang L, Wang S, Xu C, Chen X: A Comparison for Dimensionality Reduction Methods of Single-Cell RNA-seq Data. Front Genet 2021, 12:646936.

3. Ding J, Regev A: Deep generative model embedding of single-cell RNA-Seq profiles on hyperspheres and hyperbolic spaces. Nat Commun 2021, 12:2554.

4. Kobak D, Berens P: The art of using t-SNE for single-cell transcriptomics. Nat Commun 2019, 10:5416.

5. van der Maaten L, Hinton G: Visualizing Data using tSNE. J Mach Learn Res 2008, 9:2579–2605.

6. Becht E, McInnes L, Healy J, Dutertre CA, Kwok IWH, Ng LG, Ginhoux F, Newell EW: Dimensionality reduction for visualizing single-cell data using UMAP. Nat Biotechnol 2018.

7. McInnes L, Healy J, Melville J: UMAP: Uniform Manifold Approximation and Projection for Dimension Reduction. arXiv 2018.

8. Hao Y, Hao S, Andersen-Nissen E, Mauck WM, 3rd, Zheng S, Butler A, Lee MJ, Wilk AJ, Darby C, Zager M, et al: Integrated analysis of multimodal single-cell data. Cell 2021, 184:3573–3587 e3529.

9. Wolf FA, Angerer P, Theis FJ: SCANPY: large-scale single-cell gene expression data analysis. Genome Biol 2018, 19:15.

10. Dou J, Liang S, Mohanty V, Miao Q, Huang Y, Liang Q, Cheng X, Kim S, Choi J, Li Y, et al: Bi-order multimodal integration of single-cell data. Genome Biol 2022, 23:112.

11. Hie B, Bryson B, Berger B: Efficient integration of heterogeneous single-cell transcriptomes using Scanorama. Nat Biotechnol 2019, 37:685–691.

12. Peyvandipour A, Shafi A, Saberian N, Draghici S: Identification of cell types from single cell data using stable clustering. Sci Rep 2020, 10:12349.

13. Szabo PA, Levian HM, Miron M, Snyder ME, Senda T, Yuan J, Cheng YL, Bush EC, Dogra P, Thapa P, Farber DL, Sims PA: Single-cell transcriptomics of human T cells reveals tissue and activation signatures in health and disease. Nat Commun 2019, 10:4706.

14. Saelens W, Cannoodt R, Todorov H, Saeys Y: A comparison of single-cell trajectory inference methods. Nat Biotechnol 2019, 37:547–554.

15. Cao J, Spielmann M, Qiu X, Huang X, Ibrahim DM, Hill AJ, Zhang F, Mundlos S, Chrisaansen L, Steemers FJ, Trapnell C, Shendure J: The single-cell transcriptional landscape of mammalian organogenesis. Nature 2019, 566:496–502.

16. Wang X, Yang L, Wang YC, Xu ZR, Feng Y, Zhang J, Wang Y, Xu CR: Comparative analysis of cell lineage differentiation during hepatogenesis in humans and mice at the single-cell transcriptome level. Cell Res 2020, 30:1109–1126.

17. Liu X, Ouyang JF, Rossello FJ, Tan JP, Davidson KC, Valdes DS, Schroder J, Sun YBY, Chen J, Knaupp AS, et al: Reprogramming roadmap reveals route to human induced trophoblast stem cells. Nature 2020, 586:101–107.

18. Chari T, Pachter L: The specious art of single-cell genomics. PLoS Comput Biol 2023, 19:e1011288.

19. Heiser CN, Lau KS: A Quantitative Framework for Evaluating Single-Cell Data Structure Preservation by Dimensionality Reduction Techniques. Cell Rep 2020, 31:107576.

20. Kobak D, Linderman GC: Initialization is critical for preserving global data structure in both t-SNE and UMAP. Nat Biotechnol 2021, 39:156–157.

21. Alquicira-Hernandez J, Powell JE, Phan TG: No evidence that plasmablasts transdifferentiate into developing neutrophils in severe COVID-19 disease. Clin Transl Immunology 2021, 10:e1308.

22. Cooley SM, Hamilton T, Aragones SD, Ray JCJ, Deeds EJ: A novel metric reveals previously unrecognized distortion in dimensionality reduction. bioRxiv 2019.

23. Aizarani N, Saviano A, Sagar, Mailly L, Durand S, Herman JS, Pessaux P, Baumert TF, Grun D: A human liver cell atlas reveals heterogeneity and epithelial progenitors. Nature 2019, 572:199–204.

24. Ramachandran P, Dobie R, Wilson-Kanamori JR, Dora EF, Henderson BEP, Luu NT, Portman JR, Matched KP, Brice M, Marwick JA, et al: Resolving the fibrotic niche of human liver cirrhosis at single-cell level. Nature 2019, 575:512–518.

25. Zhang M, Yang H, Wan L, Wang Z, Wang H, Ge C, Liu Y, Hao Y, Zhang D, Shi G, et al: Single-cell transcriptomic architecture and intercellular crosstalk of human intrahepatic cholangiocarcinoma. J Hepatol 2020, 73:1118–1130.

26. MacParland SA, Liu JC, Ma XZ, Innes BT, Bartczak AM, Gage BK, Manuel J, Khuu N, Echeverri J, Linares I, et al: Single cell RNA sequencing of human liver reveals distinct intrahepatic macrophage populations. Nat Commun 2018, 9:4383.

27. Payen VL, Lavergne A, Alevra Sarika N, Colonval M, Karim L, Deckers M, Najimi M, Coppieters W, Charloteaux B, Sokal EM, El Taghdouini A: Single-cell RNA sequencing of human liver reveals hepatic stellate cell heterogeneity. JHEP Rep 2021, 3:100278.

28. Wilk AJ, Rustagi A, Zhao NQ, Roque J, Maranez-Colon GJ, McKechnie JL, Ivison GT, Ranganath T, Vergara R, Hollis T, et al: A single-cell atlas of the peripheral immune response in patients with severe COVID-19. Nat Med 2020, 26:1070–1076.

29. Xia L, Lee C, Li JJ: scDEED: a statistical method for detecting dubious 2D single-cell embeddings and optimizing t-SNE and UMAP hyperparameters. bioRxiv 2023.

30. Chen EY, Tan CM, Kou Y, Duan Q, Wang Z, Meirelles GV, Clark NR, Ma’ayan A: Enrichr: interactive and collaborative HTML5 gene list enrichment analysis tool. BMC BioinformaGcs 2013, 14:128.

31. Kuleshov MV, Jones MR, Rouillard AD, Fernandez NF, Duan Q, Wang Z, Koplev S, Jenkins SL, Jagodnik KM, Lachmann A, et al: Enrichr: a comprehensive gene set enrichment analysis web server 2016 update. Nucleic Acids Res 2016, 44:W90–97.

32. Xie Z, Bailey A, Kuleshov MV, Clarke DJB, Evangelista JE, Jenkins SL, Lachmann A, Wojciechowicz ML, Kropiwnicki E, Jagodnik KM, Jeon M, Ma’ayan A: Gene Set Knowledge Discovery with Enrichr. Curr Protoc 2021, 1:e90.

33. Gillespie M, Jassal B, Stephan R, Milacic M, Rothfels K, Senff-Ribeiro A, Griss J, Sevilla C, Madhews L, Gong C, et al: The reactome pathway knowledgebase 2022. Nucleic Acids Res 2022, 50:D687–D692.

