## Supplemental figures for "cellstruct: Metrics scores to quantify the biological preservation between two embeddings"

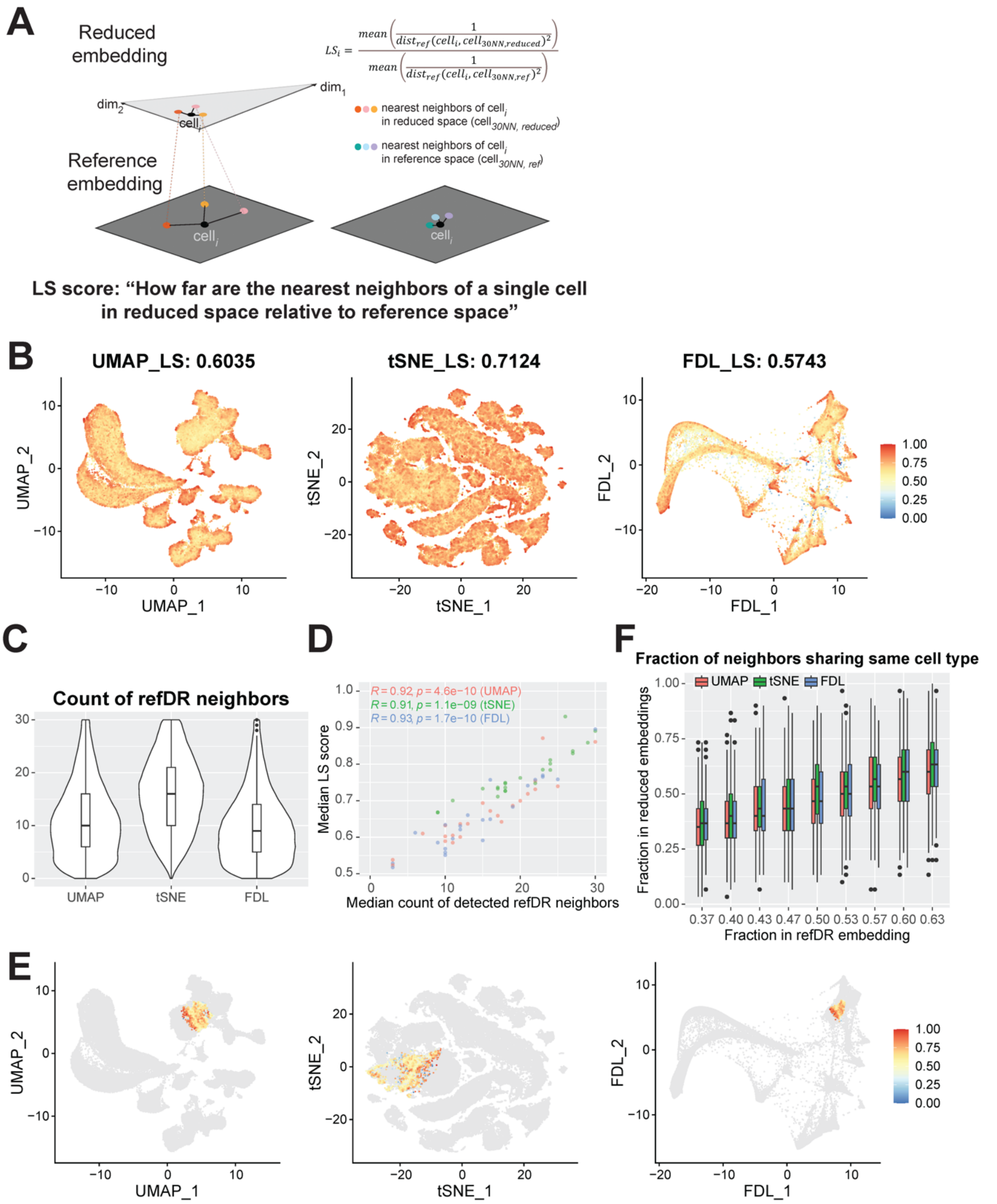


$$LS_i = \frac{mean\left(\frac{1}{dist_{ref}(cell_i, cell_{30NN,reduced})^2}\right)}{mean\left(\frac{1}{dist_{ref}(cell_i, cell_{30NN,ref})^2}\right)}$$

**Figure S1.** Evaluation of local cell-cell relationships via local single-cell (LS) metric in human liver cell atlas. (A) Schematic showing the calculation of LS metric. LS score is defined as the ratio between the average reciprocal-squared-distance of the 30 nearest neighboring (NN) cells in reduced embedding and that of the 30NN cells in reference embedding. (B) The LS metric was calculated for UMAP, t-SNE, and FDL embeddings respectively, and the average scores were indicated in the title. (C) Violin plots exhibiting the number of overlap between the 30NN cells in different reduced embeddings and the 150 NN cells in refDR (i.e. reference) embedding. (D) Scatter plot showing the correlation between the median count of overlapping NN cells and the median LS scores for each cell type in different reduced embeddings, with the correlations and *p*-values indicated at the top left corner. (E-F) 2,189 “boundary” cells between CD4 and CD8 T cells were selected in the refDR embedding. These “boundary” cells have only CD4 and CD8 T cells in their 30NN cells in the refDR embedding, out of which 11-19 cells originate from either cell type. The fraction of 30 NN cells, from respective embedding, sharing the same cell type label as the “boundary” cells was illustrated in (E). These fractions of reduced NN cells were compared against those of reference NN cells using violin plots (F).

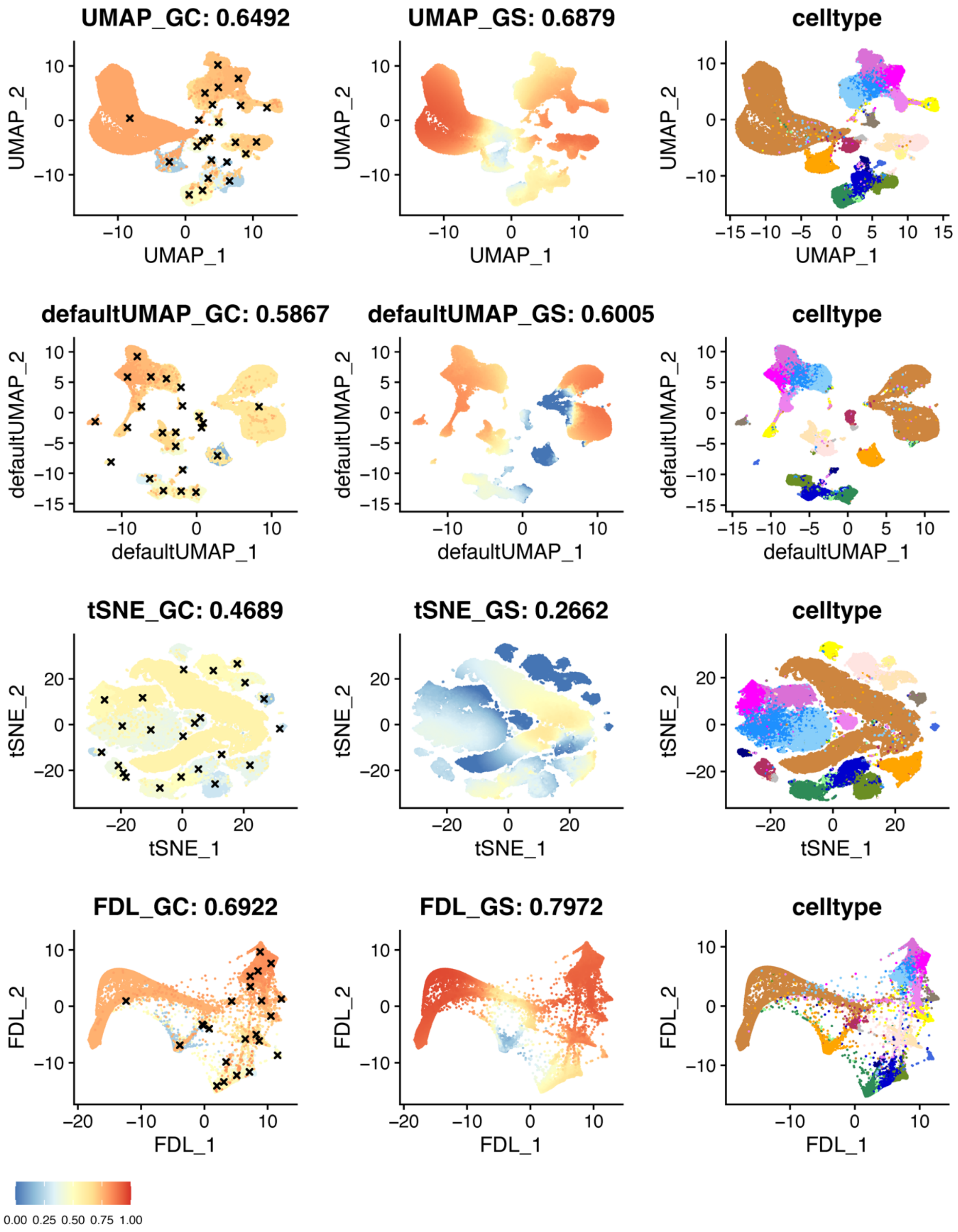


**Figure S2.** An example of cellstruct output using the human liver cell atlas. The GC (left column) and GS (middle column) scores were calculated for four different reduced embeddings: UMAP (tuned parameters: Euclidean metric, 50 n.neighbors, and 0.5 min.dist), UMAP (default parameters: cosine metric, 30 n.neighbors, and 0.3 min.dist), t-SNE, and FDL, with refDR as the reference embedding. The mean of these scores was shown in the title of each panel. The annotated cell types (right column) were shown for each embedding with the same color scheme depicted as in Figure 1C.

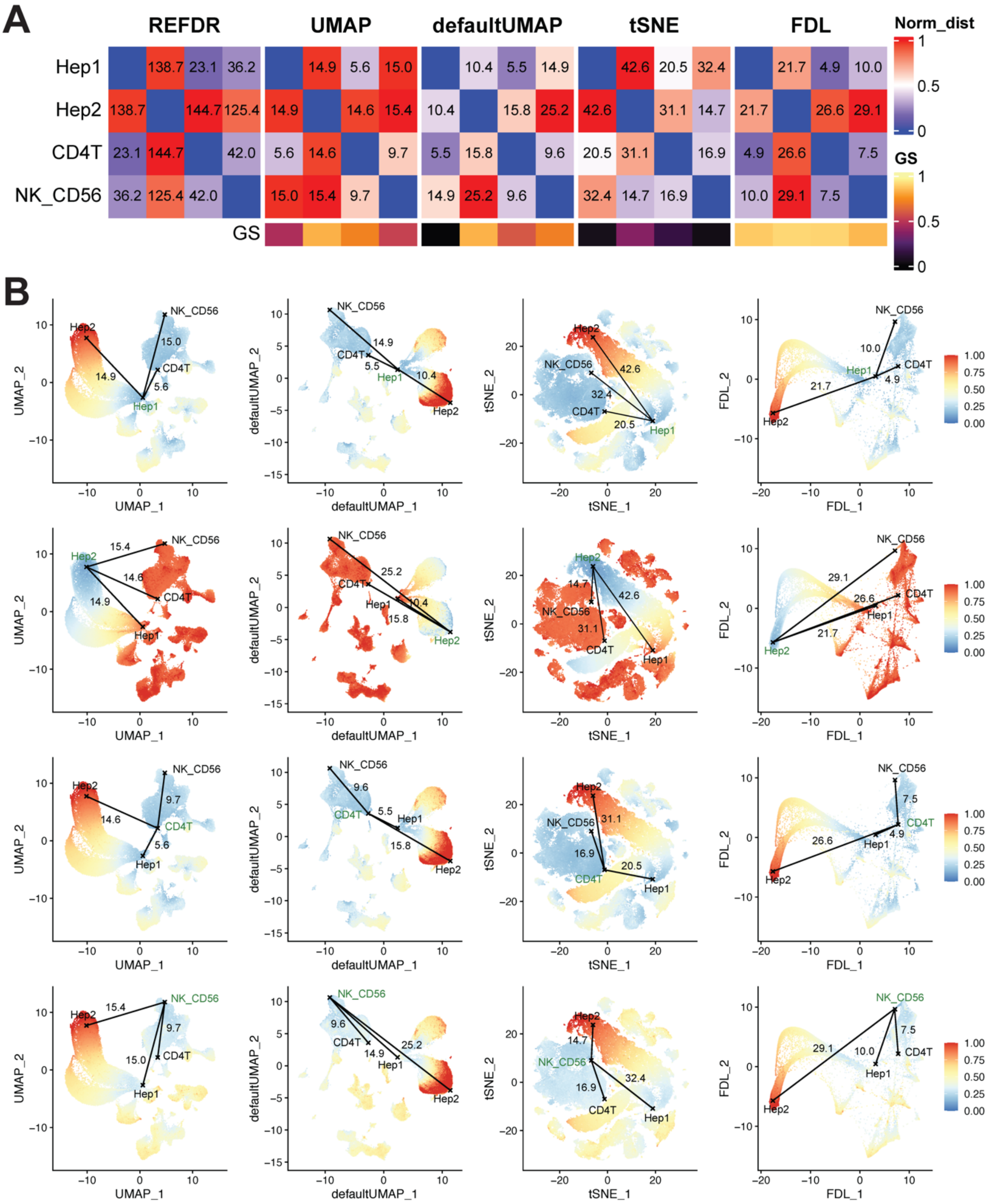


**Figure S3.** Four single cells were selected as an illustration for the comparison between different reduced embeddings. These cells are comprised of two hepatocytes: Hep1 and Hep2, one CD4 T, and one NK_CD56bright (labeled as NK_CD56). (A) Five heatmaps were colored by the pairwise distances in the reference embedding (refDR) and four different reduced embeddings respectively, normalized by the maximum distance in each distance matrix, and they were labeled with the exact pairwise distances. The GS scores of these four cells were colored at the bottom of heatmaps for each reduced embedding. (B) The normalized pairwise reference distances between each of the four cells (labeled green) and the remaining cells, were colored in the respective reduced projections, with the exact pairwise reduced distances among these four cells indicated as text in each panel. E.g. The top row represents the pairwise distances between Hep1 and the remaining cells in UMAP, default UMAP, t-SNE, and FDL projections.

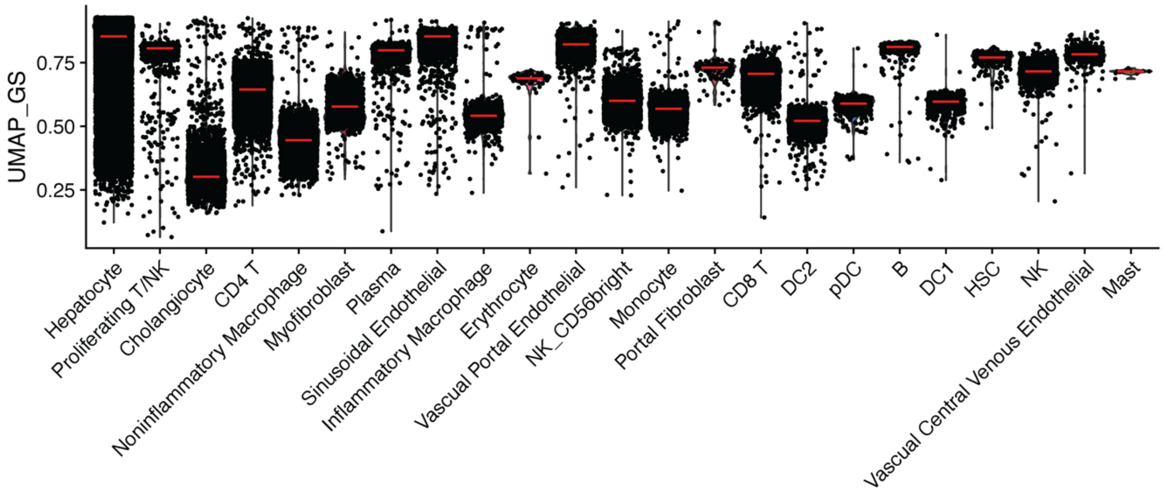


**Figure S4**. Distribution of GS scores for UMAP embedding in human liver cell atlas. Violin plots showing the GS scores distribution in each cell type, ordered by decreasing variance in the cell type's GS scores. The red line indicates the median GS score for each cell type.

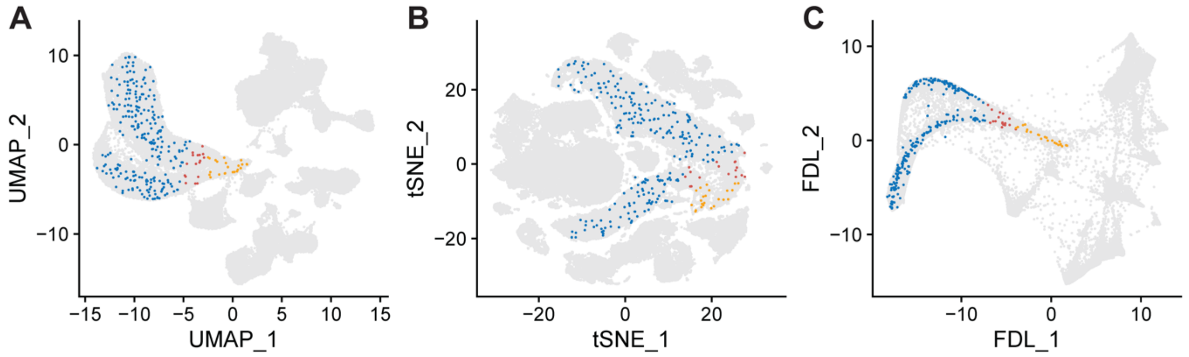


**Figure S5.** Dimension reductions showing three groups of hepatocytes. These three groups, identified in the 284 hepatocytes (Figure 1F), were illustrated in (A) UMAP, (B) t-SNE, and (C) FDL embeddings respectively.

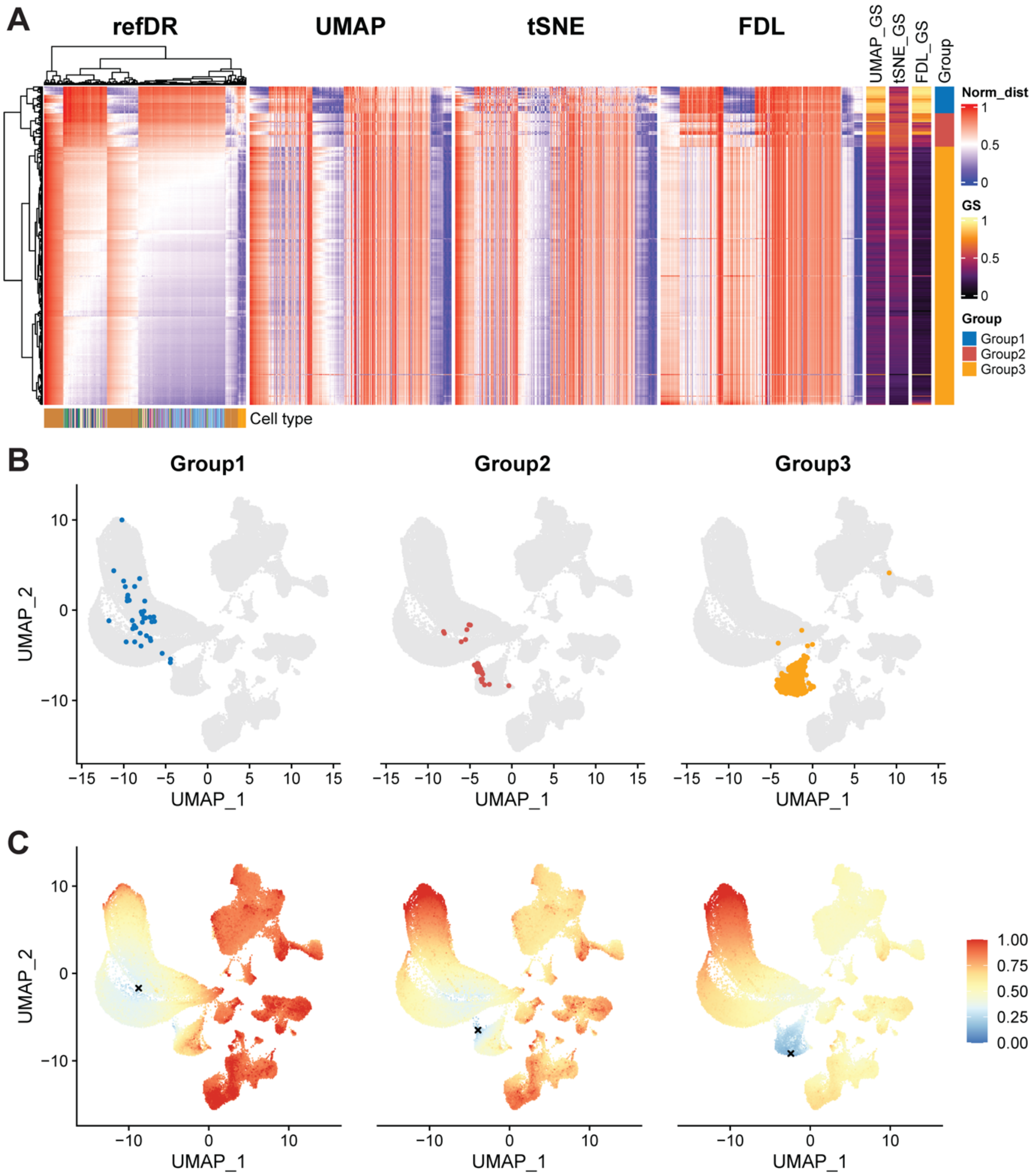


**Figure S6**. Investigating cholangiocytes in the human liver cell atlas using cellstruct. (A) Heatmaps illustrating the normalized cell-cell distances between 428 cholangiocytes (refer to Methods for the selection of these cholangiocytes) and 1,000 waypoint cells in refDR (i.e. reference) and reduced embeddings, along with the calculated GS scores for each reduced embedding: UMAP, t-SNE, and FDL. Three groups of cells were identified based on the clustering pattern detected in the normalized refDR distances (leftmost heatmap). Amongst these three groups, 81 "outlier" cells (from Groups A and B) were detected. (B) The three groups of cholangiocytes were shown in the UMAP embeddings. (C) One of the cells was selected from each group, and the UMAPs were colored by the normalized refDR distances between the selected (marked by 'X') cell and all other cells.

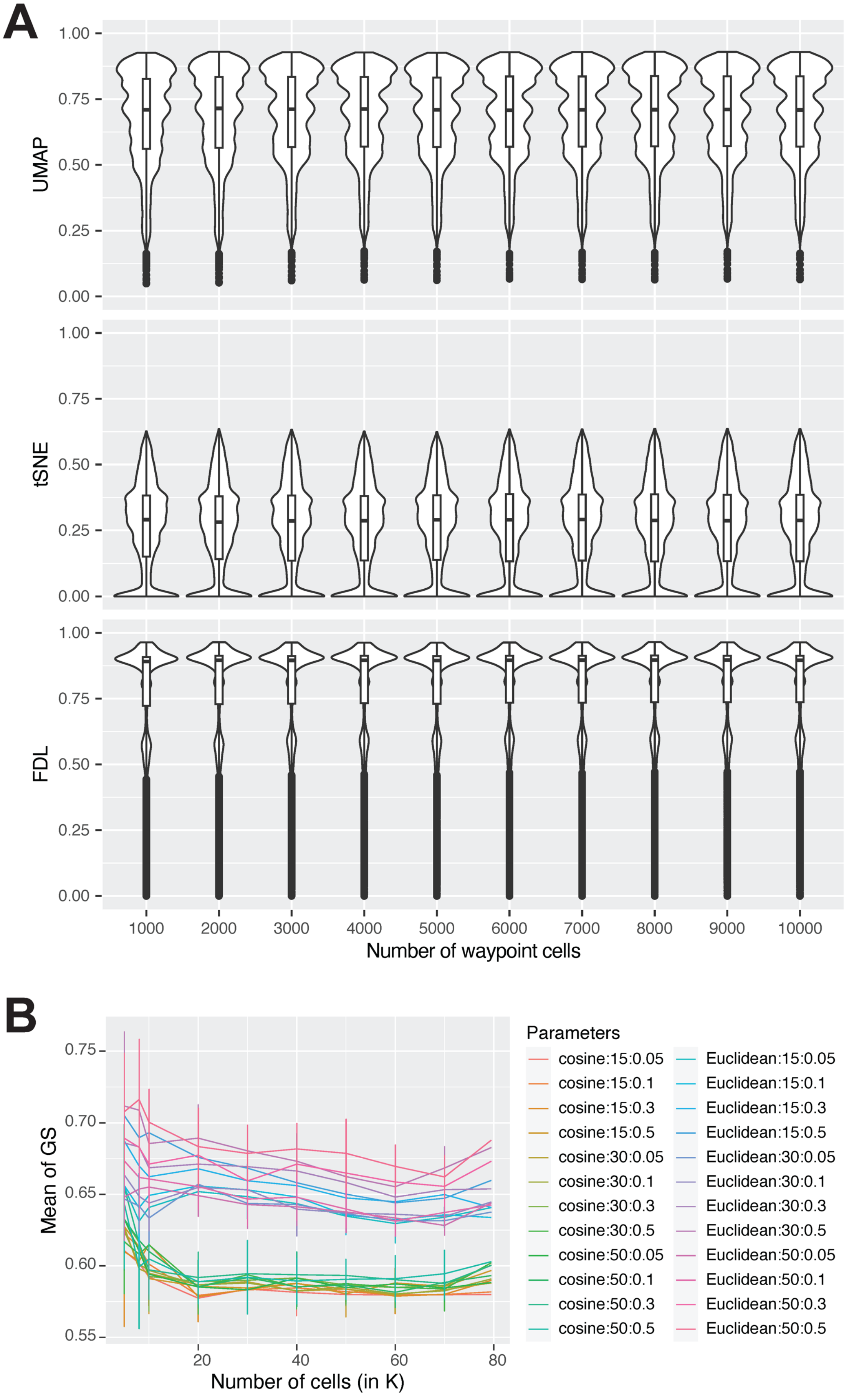
A
UMAP
tSNE
FDL
1.00
0.75
0.50
0.25
0.00
1000
2000
3000
4000
5000
6000
7000
8000
9000
10000
Number of waypoint cells
B
Mean of GS
0.75
0.70
0.65
0.60
0.55
20
40
60
80
Number of cells (in K)
Parameters
cosine:15:0.05
cosine:15:0.1
cosine:15:0.3
cosine:15:0.5
cosine:30:0.05
cosine:30:0.1
cosine:30:0.3
cosine:30:0.5
cosine:50:0.05
cosine:50:0.1
cosine:50:0.3
cosine:50:0.5
Euclidean:15:0.05
Euclidean:15:0.1
Euclidean:15:0.3
Euclidean:15:0.5
Euclidean:30:0.05
Euclidean:30:0.1
Euclidean:30:0.3
Euclidean:30:0.5
Euclidean:50:0.05
Euclidean:50:0.1
Euclidean:50:0.3
Euclidean:50:0.5

**Figure S7.** Stability analysis of GS metric using human liver cell atlas. (A) The number of waypoint cells was varied between 1K to 10K for UMAP, t-SNE and FDL embeddings respectively (top to bottom panels). The median GS score was illustrated for all these sets in the violin plots. (B) The number of cells was downsampled to 5K, 8K, 10K, 20K, …, 60K, and 70K for 10 times respectively. The GS scores were computed for each cell in these subsampled sets, with the UMAP embedding of each set being re-generated under 24 different combinations of hyperparameters: metric (cosine or Euclidean), n.neigbors (15, 30, and 50), and min.dist (0.05, 0.1, 0.3, and 0.5). These GS scores were then averaged across a single UMAP embedding set, and the distribution of the mean GS score was plotted against the number of cells as above, with the complete dataset as the final data point.

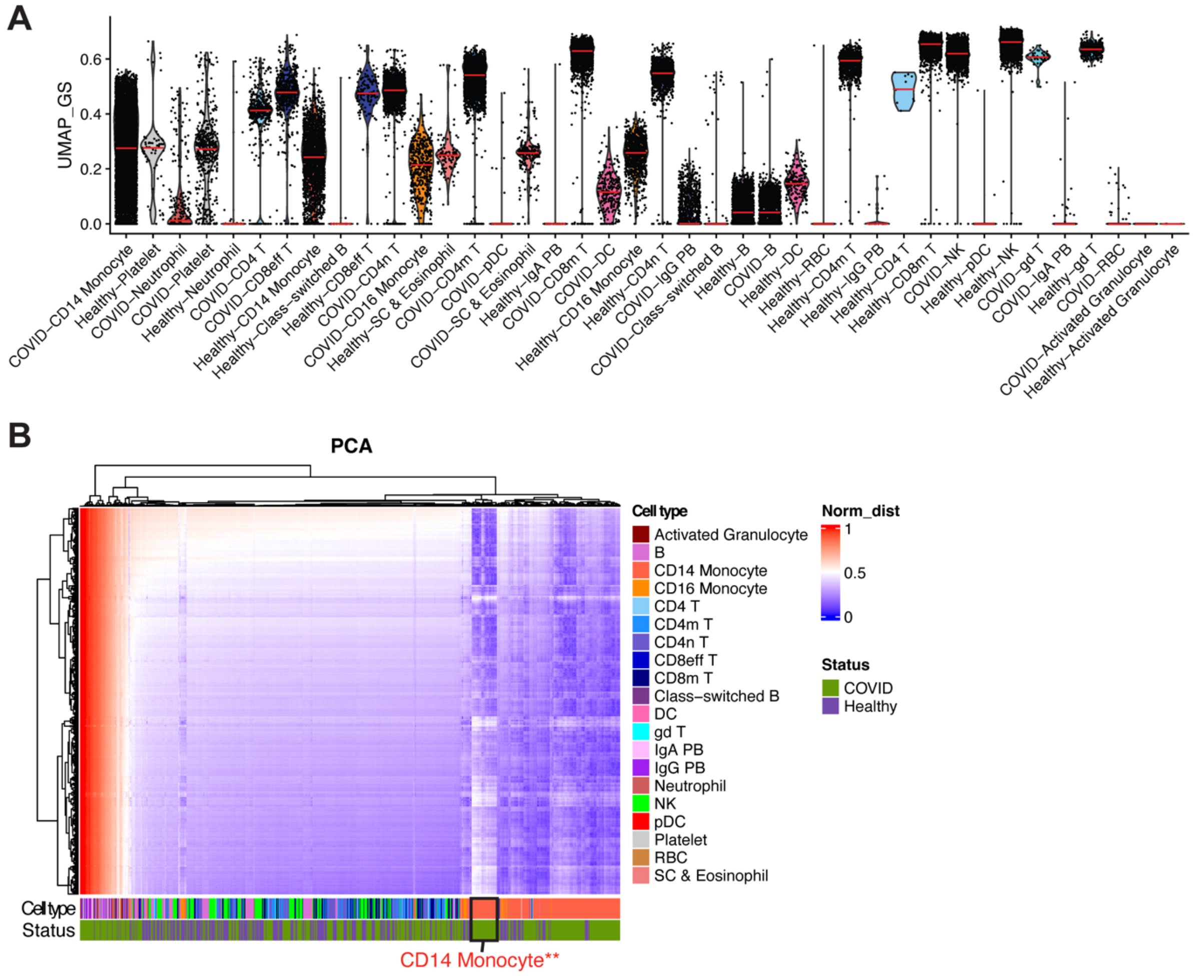


**Figure S8.** Applying cellstruct to the COVID peripheral immune atlas. (A) Violin plots portraying the distribution of GS scores for UMAP embedding in each COVID/healthy cell type, ordered by decreasing variance of GS scores. The red line indicates the median GS score for each cell type. (B) Heatmap representing the normalized cell-cell distances between the 20% of randomly sampled COVID-CD14 Monocytes (1,657) and 1,000 waypoint cells in PCA (i.e. reference) embedding, with the cell type and COVID/healthy status of 1,000 waypoint cells annotated at the bottom of heatmap. The black box marked a subgroup of COVID-CD14 Monocytes that did not cluster with the remaining COVID-CD14 Monocytes.

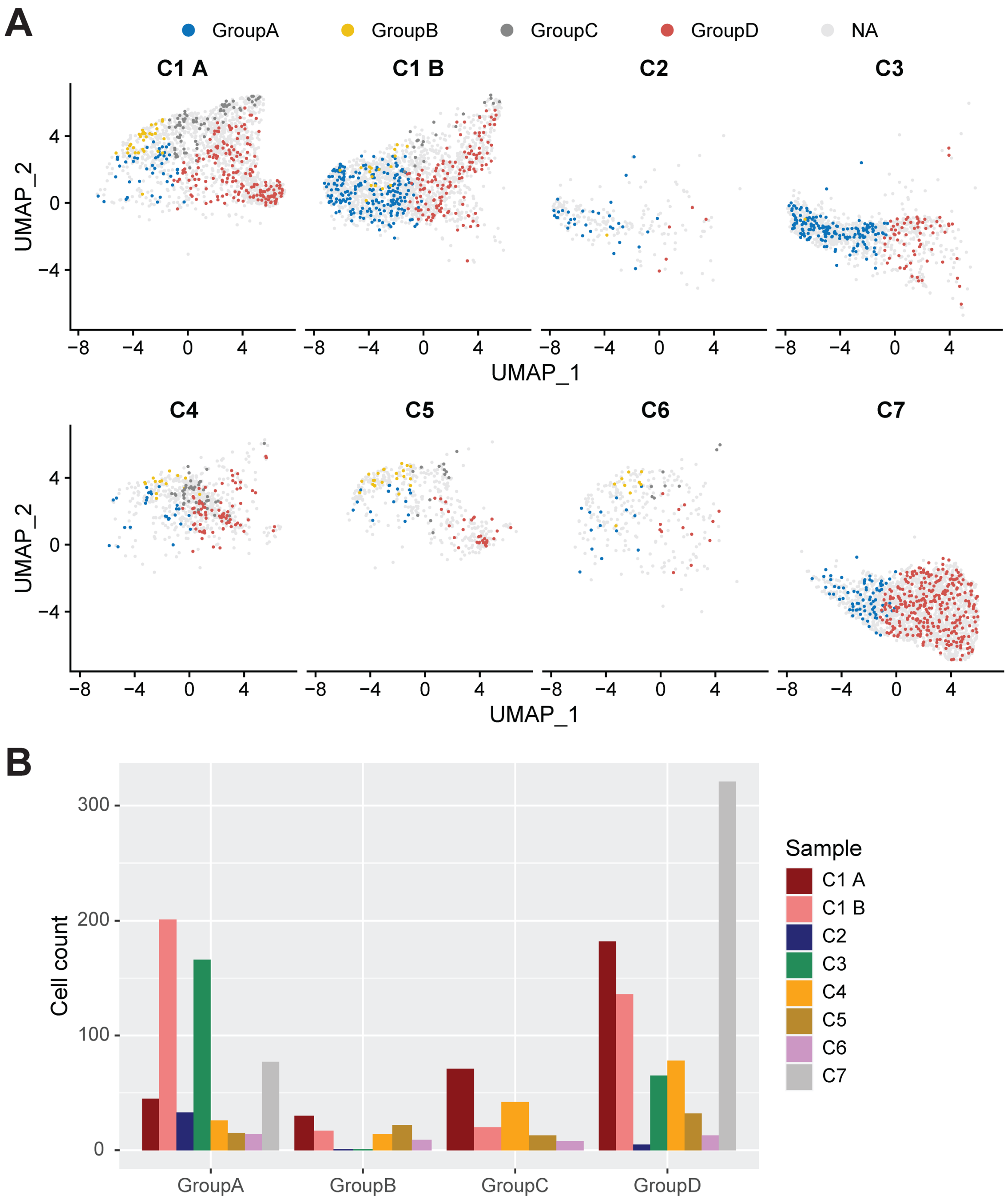


**Figure S9**. Corroboration of COVID-CD14 Monocyte groups, identified by cellstruct, with the author's original analysis. (The analysis is explained in the original publication's Figure 2, particularly in Figure 2h). The location of four groups was illustrated on the UMAP embeddings of each sample in (A). There were very few Groups B and C cells (enriched in interferon signaling) detected in patients C2 (1), C3 (1), and C7(0). The number of cells detected from each sample in each monocyte group was tabulated in Table S2 and was shown in the bar plot (B).

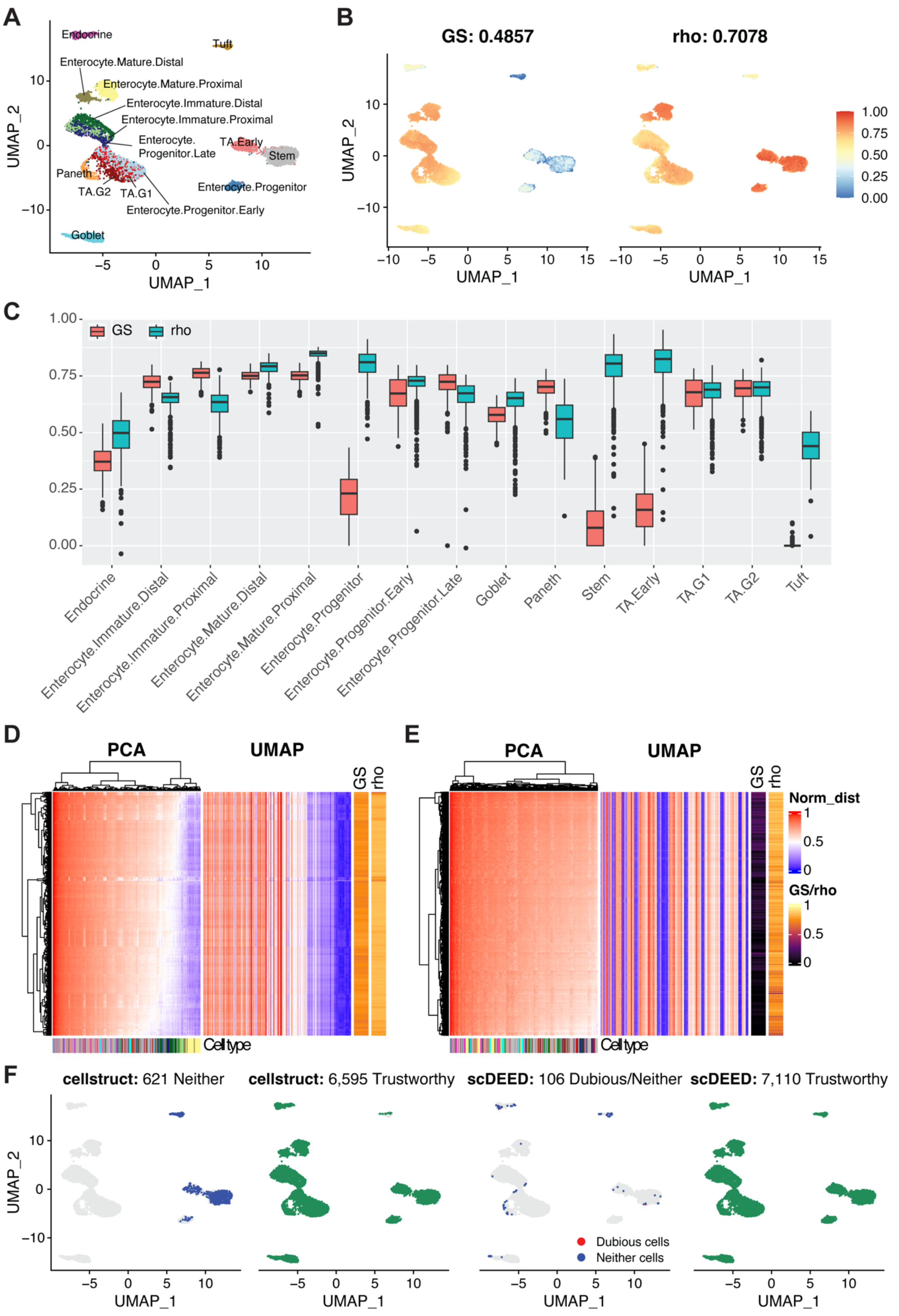


**Figure S10.** Comparative analysis between scDEED and cellstruct using Simulated Dataset 1. (A) UMAP embedding of simulated dataset was colored by annotated cell types. (B) The GS and rho scores, calculated by cellstruct and scDEED respectively, were illustrated on the UMAP projections, with their mean indicated in the title. (C) Boxplots of GS and rho scores from each cell type. 581 Enterocytes.Mature.Proximal and 1,267 Stem cells were selected to show the normalized cell-cell distances between these cells and 1,000 waypoint cells in both PCA and UMAP embeddings, elucidating the high GS and high rho values in (D, Enterocytes.Mature.Proximal) and low GS and high rho values in (E, Stem). (F) UMAP embeddings showing dubious, neither (i.e. neither dubious nor trustworthy), and trustworthy cells determined using either cellstruct's GS metric or scDEED's rho score. Using the similar approach as scDEED, the GS scores were compared against a null distribution of GS scores generated from a permuted object provided by scDEED authors. Cellstruct identified 6,595 trustworthy and 621 neither cells, while scDEED determined 7,110 trustworthy, 2 dubious, and 104 neither cells.

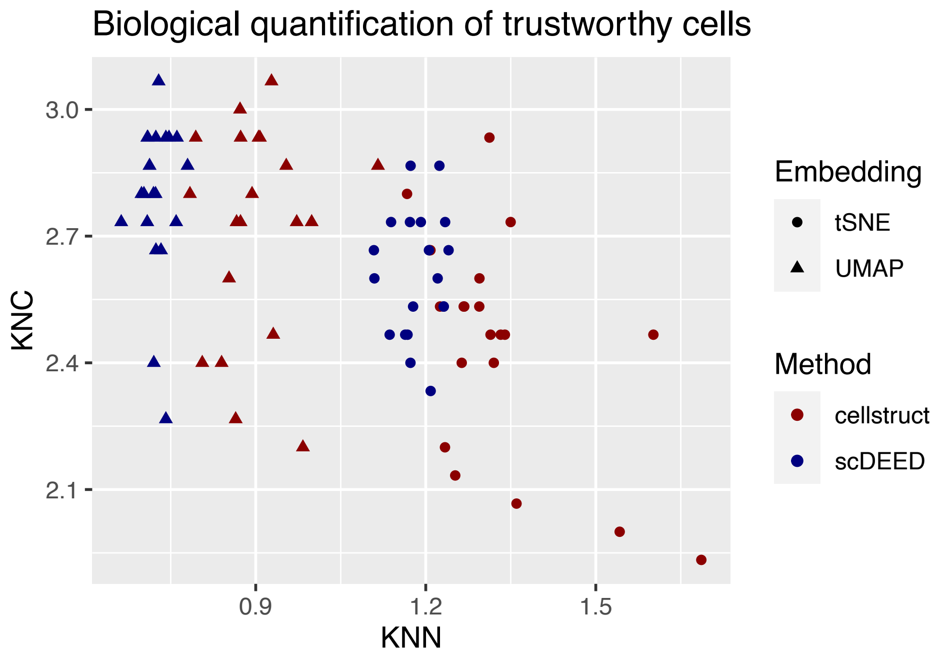


**Figure S11.** Scatterplot of KNC and KNN values of t-SNE and UMAP embeddings in 20 simulated datasets. These values were calculated from the trustworthy cells, identified by cellstruct and scDEED respectively. Using paired t-test, cellstruct showed significantly higher KNN values in both t-SNE (mean difference:0.15, $p$:2.29x10$^{-4}$) and UMAP (mean difference:0.18, $p$:3.68x10$^{-9}$) embeddings, while scDEED had higher KNC values in t-SNE (mean difference:0.17, $p$:0.01) but not in UMAP (mean difference:0.07 higher in scDEED, $p$:0.35) embedding.
