## Supplemental text for "cellstruct: Metrics scores to quantify the biological preservation between two embeddings"

### Assessing local cell-cell relationships via local single-cell (LS) metric in human liver cell atlas

As part of the cellstruct package, we implemented three metrics assessing the preservation of local or global relationships between a reduced (e.g. UMAP) and a reference (e.g. PCA) embedding. For local cell-cell relationships, we devised the LS metric, which quantifies whether the 30 nearest neighboring (NN) cells in the reduced embedding remain close to the single cell of interest in the reference embedding. To achieve this, we calculated the ratio of the relative distances of these 30NN cells in reduced embedding to the distances of the 30NN cells in reference embedding (refer to the formula in Methods section). Note that all distances were computed in the reference space (Figure S1A). These LS scores were calculated for each single cell, for respective embeddings: UMAP (after tuning), t-SNE, and FDL. t-SNE embedding showed the highest mean LS score across all cells, corroborating with the optimization objective of t-SNE, which prioritizes local structure of data. Furthermore, we observed a phenomenon, in which cells lining the edges of a cluster i.e. edge cells generally have higher LS scores, indicating that these cells' reduced neighbors (i.e. nearest neighbors in reduced embedding) remain close in the reference space. This is probably because the 30NN cells of these edge cells are constrained in the space where they can be distributed as compared to the non-edge cells that were located further away from the edges (Figure S1B).

In addition to the LS metric, we counted the overlap between the 30NN cells in reduced embedding and the 150NN cells in reference embedding. Similarly, t-SNE showed the highest median number of overlapping NN cells (t-SNE:16, UMAP:10, FDL: 9), corroborating the results from the LS metric. When grouping the single cells by cell type, the median count of these overlapping neighbors is significantly correlated with the median LS scores ($r$>0.9, $p$<1e-8) (Figures S1C-D).

Based on the LS metric and overlapping NN cells, t-SNE embedding was shown to be the best embedding (as compared to UMAP and FDL) in preserving the local structure. Therefore, we were interested to study if highly related cell types are better separated in t-SNE embedding. Using CD4 and CD8 T cells as an illustration, 2,189 "boundary" cells between CD4 and CD8T were identified from the reference embedding. These "boundary" cells were selected such that they have only CD4 and CD8 T cells in their 30NN cells in the reference embedding, out of which 11-19 cells originate from either cell type. Next, using the t-SNE embedding, we calculated the fraction of 30NN cells sharing the same cell type label as these "boundary" cells. This was then repeated for the UMAP and FDL embeddings, and we observed no significant difference among t-SNE, UMAP, and FDL when compared to the reference embedding (Figures S1E-F). Although t-SNE embedding better preserves local cell-cell relationships in terms of the NN cells, this property is not helpful in separating the transcriptionally similar cell types like CD4/CD8 T cells. Thus, we choose to omit this metric and instead focus on the global single-cell (GS) and global cluster (GC) scores in the main text.

### Tuning UMAP hyperparameters to improve GS scores

From the UMAP embedding calculated using Seurat's default hyperparameters (cosine metric, 30 n.neighbors, and 0.3 min.dist), we observed that the GS scores within hepatocytes are highly non-uniform, and most cholangiocytes were assigned with low GS scores. This prompted us to vary the UMAP hyperparameters, considering various combinations of them (cosine or Euclidean metric; n.neighbors: 15, 30, and 50; min.dist: 0.05, 0.1, 0.3, and 0.5), with the goal of improving the GS scores. The embedding, which has the highest mean GS score, was selected and termed tuned UMAP (this was implemented using the tuneUMAP function in our cellstruct package). The mean GS score from the tuned UMAP is higher than that of default UMAP (0.69 vs 0.60). We also noticed that the hepatocytes and cholangiocytes (and part of monocytes) have higher GS scores after tuning by being positioned in a more "accurate" global position (Figure S2). The stability of tuneUMAP function was evaluated for this dataset, by downsampling the number of cells, with each downsampling being performed 10 times respectively. Each of these downsampled datasets was assessed for their GS scores under various combinations of UMAP hyperparameters. Overall, the mean GS score is fairly consistent beyond 20K cells. Also, UMAPs generated using the Euclidean metric tend to produce higher mean GS scores as compared with those with the cosine metric (Figure S7B).

**Comparative analysis of reduced embeddings using human liver cell atlas**
As discussed in the main text, different embeddings (i.e. UMAP, t-SNE, and FDL) have different GS scores, reflecting different degree of preservation of the global cell-cell relationships. To further visualize these differences in the GS scores, four single cells (two hepatocytes, one CD4 T, and one NK_CD56bright [labeled as NK_CD56]), which were positioned at the different ends of the hepatocyte or lymphoid "islands" on the default UMAP were selected. Their default UMAP distances are relatively similar (Hep1-Hep2: 10.4 and CD4T-NK_CD56: 9.6), but the distribution of GS scores across T/NK group is much more uniform than those of hepatocytes (Figure S2). Therefore, we were intrigued to study these distances in different embeddings, by illustrating the pairwise distances among these cells (Figure S3A) and between each of these cells and all other cells (Figure S3B, colored by normalized reference distances in different reduced embeddings). The pairwise distances among these cells in the reference embedding highlighted the large distance between Hep2 and the other three cells, implying the transcriptomic dissimilarity of this Hep2 with other cells, including Hep1. Similarly, this was demonstrated in the different reduced embeddings, where hepatocytes near the "lobes" of UMAP (i.e. location of Hep2) are clearly different from all other cells, and this difference is captured in all embeddings (Figure S3B). FDL is the best reduced embedding in terms of recapitulating this dissimilarity and the relationship with the other cells, providing high GS scores to all these four cells. The tuned UMAP performed better than default UMAP, by positioning Hep2 further away from Hep1 (i.e. elongating the "lobes" of hepatocytes, which was also observed in the FDL embedding), and this improved the GS score of Hep1. Nevertheless, the tuned UMAP distance between Hep1 and Hep2 is similar to those of Hep1 and NK_CD56, which failed to represent the underlying distances in the reference embedding, where Hep2 is more dissimilar to Hep1 as compared to NK_CD56. As for the t-SNE embedding, although it showed the largest distance between Hep1 and Hep2, Hep2 became the closest cell to NK_CD56 as compared to CD4T, which is not biologically correct (Figure S3). Overall, t-SNE embedding gives a poor representation of the global cell-cell relationships in the liver cell atlas data, as depicted by the low GS scores across all cells (Figure S2).

**Comparative analysis between cellstruct and scDEED using simulated dataset1**
Simulated data from Xia et al., was generated from an experimentally derived scRNA-seq dataset of mouse small intestinal epithelial cells (10,000 genes and 7,217 cells),

comprising of enterocytes, tuft, Paneth, stem, goblet, and transit-amplifying (TA) cells [1]. Both cellstruct (GS) and scDEED (rho) scores were utilized to assess the reliability of UMAP embedding. Both scores have similar trends across cell type, with the rho score generally depicting higher values, except in tuft, stem, TA.early, and Enterocyte.Progenitor cells. Hence, we selected the Enterocyte.Mature.Proximal (high GS and high rho) and Stem (low GS and high rho) cells for visualization of the normalized pairwise distances between these cells and 1,000 waypoint cells in the reference PCA and UMAP embeddings respectively. We observed concordance between the PCA and UMAP distances for the Enterocyte.Mature.Proximal cells but not Stem cells, suggesting that the correlation between PCA and UMAP embeddings should be low for Stem cells, as confirmed by the GS values (Figures S10A-E).

Following the same approach as scDEED, we classified the cells into trustworthy and dubious by comparing GS scores to a null distribution (i.e. GS scores assigned to the same permuted object generated by scDEED). Trustworthy cells are defined by their GS scores above 95 percentile of null distribution GS, whereas dubious cells are cells with their GS scores below 5 percentile of null distribution GS. For 20 simulated datasets, we detected fewer trustworthy cells using our GS score as compared to the scDEED metric, and more cells fell into “neither” range (i.e. neither statistically termed trustworthy nor dubious), as shown in Table S3. For example, in simulated dataset 1, we detected 6,595 trustworthy and 621 neither cells, while scDEED detected 7,110 trustworthy, 104 neither, and 2 dubious cells (Figure S10F).

**References**

1. Tabula Muris C, Overall c, Logistical c, Organ c, processing, Library p, sequencing, Computational data a, Cell type a, Writing g, Supplemental text writing g, Principal i: **Single-cell transcriptomics of 20 mouse organs creates a Tabula Muris.** *Nature* 2018, **562:**367-372.
